# P66 is a bacterial mimic of CD47 that binds the anti-phagocytic receptor SIRPα and facilitates macrophage evasion by *Borrelia burgdorferi*

**DOI:** 10.1101/2024.04.29.591704

**Authors:** Michal Caspi Tal, Paige S. Hansen, Haley A. Ogasawara, Qingying Feng, Regan F. Volk, Brandon Lee, Sara E. Casebeer, Grace S. Blacker, Maia Shoham, Sarah D. Galloway, Anne L. Sapiro, Beth Hayes, Laughing Bear Torrez Dulgeroff, Tal Raveh, Venkata Raveendra Pothineni, Hari-Hara SK Potula, Jayakumar Rajadas, Effie E. Bastounis, Seemay Chou, William H. Robinson, Jenifer Coburn, Irving L. Weissman, Balyn W. Zaro

## Abstract

Innate immunity, the first line of defense against pathogens, relies on efficient elimination of invading agents by phagocytes. In the co-evolution of host and pathogen, pathogens developed mechanisms to dampen and evade phagocytic clearance. Here, we report that bacterial pathogens can evade clearance by macrophages through mimicry at the mammalian anti-phagocytic “don’t eat me” signaling axis between CD47 (ligand) and SIRPα (receptor). We identified a protein, P66, on the surface of *Borrelia burgdorferi* that, like CD47, is necessary and sufficient to bind the macrophage receptor SIRPα. Expression of the gene encoding the protein is required for bacteria to bind SIRPα or a high-affinity CD47 reagent. Genetic deletion of *p66* increases phagocytosis by macrophages. Blockade of P66 during infection promotes clearance of the bacteria. This study demonstrates that mimicry of the mammalian anti-phagocytic protein CD47 by *B. burgdorferi* inhibits macrophage-mediated bacterial clearance. Such a mechanism has broad implications for understanding of host-pathogen interactions and expands the function of the established innate immune checkpoint receptor SIRPα. Moreover, this report reveals P66 as a novel therapeutic target in the treatment of Lyme Disease.

## Main Text

Cancer cells and persistently-infected cells express a variety of inhibitory ligands that dampen innate and adaptive branches of the immune system. In the case of innate immunity, phagocytic macrophages are responsible for the digestion and degradation of dead, dangerous, or foreign cells, viruses, and bacteria. Macrophage-mediated programmed cell removal is governed by pro- and anti-phagocytic ligands on mammalian cells and their cognate receptors on the surface of macrophages^1^. These ligand-receptor interactions are often referred to as “eat me” and “don’t eat me” signals, respectively. We have previously identified CD47 as a dominant “don’t eat me” signal ligand expressed by mammalian cells which engages SIRPα on the surface of macrophages^2,3^. The cytoplasmic tail of SIRPα harbors an immunoreceptor tyrosine-based inhibitory motif (ITIM), where phosphorylation events can alter downstream phagocytic potential. CD47 is a cell-surface molecule that is highly abundant on healthy red blood cells and circulating hematopoietic stem cells, among others^4^. It functions as a self-recognition ligand, allowing healthy self cells to evade innate immune-mediated phagocytosis. However, pathogen infected cells and cancer cells have been shown to hijack this pathway, upregulating CD47 expression to evade macrophage surveillance^2,3,5^. CD47-SIRPα signaling has also been implicated across other human diseases, including but not limited to atherosclerosis and neurodegenerative disease, highlighting the ubiquity and importance of this axis^6–8^.

While CD47 is well conserved across humans, its receptor SIRPα is highly polymorphic in the IgV domain where CD47 binding occurs. Notably, Barklay and Hatherley postulated that this heterogeneity may be due to pressure imposed by pathogens ^9,10^. We hypothesized that pathogenic bacteria may inhibit innate immunity through mimicry. Palm and co-workers recently reported that recombinant SIRPα binds to the surface of a strain of the commensal bacterium *Fusobacterium*^11^. We reasoned that mimics of the mammalian “don’t eat me” ligand CD47 on pathogenic bacteria would allow them to avoid a macrophage-mediated innate immune response and therefore assist in the establishment of infection.

The bacterial spirochete *Borrelia burgdorferi* (*B. burgdorferi*), the causative agent of Lyme Disease, is adept at evading innate immune clearance, which is essential to the lifestyle of this parasitic organism^12–14^. Because *B. burgdorferi* can establish a persistent infection, we hypothesized that 1) *B. burgdorferi* have a mechanism of immune evasion by inhibiting components of the human immune system beyond complement and antigenic heterogeneity produced by the *vis* locus^14,15^ and 2) *B. burgdorferi* produce a mimic of the mammalian surface protein CD47 on their surface.

We previously developed a reagent derived from SIRPα that binds human CD47 with high affinity and blocks its interaction with SIRPα, termed CV1^16^. A fusion between CV1 and the human immunoglobulin G4 heavy chain yielded a divalent, picomolar CD47-binding reagent termed CV1-G4 that binds human CD47 50,000-fold more strongly than SIRPα. To determine whether *B. burgdorferi* express a cell surface protein with structural similarity to CD47, we performed flow cytometry experiments. Specifically, we tested whether CV1-G4 and a commercially-available anti-CD47 antibody raised against a single epitope MIAP410 would bind to the surface of *B. burgdorferi* strain B31-A3. We determined that CV1-G4 binds to the surface of *B. burgdorferi* strain B31-A3 well above isotype-control or MIAP410 staining (Figure 1A).

**Figure 1.**
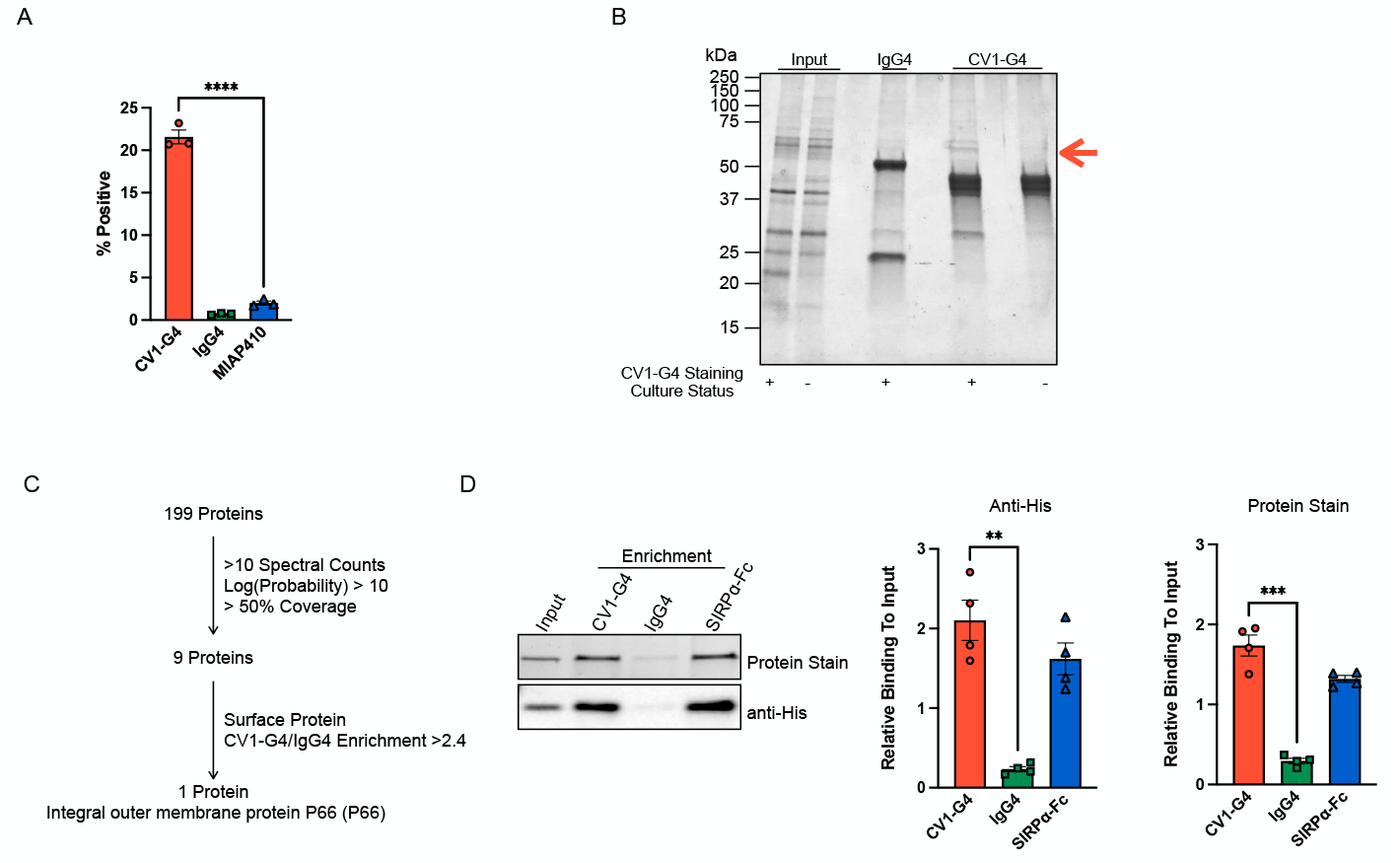
The surface protein P66 binds to a high-affinity CD47 binding reagent and the CD47 receptor SIRPα. A. *B. burgdorferi* stain for the high-affinity CD47 blocking reagent CV1-G4 by flow cytometry analysis. **** p-value < 0.0001, df = 4, N = 4; Unpaired 2-tailed t-test. B. *B. burgdorferi* was cultured under conditions to stain either positively or negatively for CV1-G4 as confirmed by flow cytometry. Bacteria were lysed under non-denaturing conditions, and lysate was subjected to enrichment with CV1-G4 or IgG4 prior to SDS-PAGE. Gel bands of interest were excised and subjected to in-gel trypsin digestion followed by mass spectrometry analysis. Arrow denotes putative P66 band. C. Data filtration parameters identified P66 as a putative CV1-G4 binding protein. D. Recombinant his-tagged P66 is enriched by CV1-G4 or SIRPα but not isotype control as determined by *in vitro* binding assay. A representative blot is shown as is quantification from 4 replicates. ** p-value = 0.0065, N = 4, df = 3. Paired two-tailed t-test; *** p-value < 0.0008, N = 4, df = 3;. Error bars calculated as SEM.

We next sought to identify the protein(s) recognized by our CD47 probe by mass spectrometry (MS). Immunoprecipitation experiments were performed from *B. burgdorferi* strain B31-A3 cultures staining positive or negative for CV1-G4 binding as determined by flow cytometry. Enriched proteins were separated by SDS-PAGE and stained with colloidal blue (Figure 1B). Gel bands of interest which appeared visibly enriched in the CV1-G4-positive lysate were excised across all three experimental conditions (CV1-G4-positive CV1-G4 enriched; CV1-G4 positive IgG4 enriched, CV1-G4-negative CV1-G4 enriched) and analyzed by MS. P66 was identified as the highest-confidence target for CV1-G4 binding, taking into consideration cellular localization and relative enrichment (Figure 1C and Supplemental Table 1).

P66 is a known antigen in *B. burgdorferi* infection^17^. In our prior murine infection model, *p66-*deficient *B. burgdorferi* (background strain B31-A3) are cleared from the site of infection within 48h, prior to the timeline of an adaptive immune response^18^. Furthermore, mice inoculated with these bacteria failed to launch an antibody-mediated response. Like CD47, P66 is an integrin binding protein, but the deficiency in persistent infectivity for *Δp66* bacteria cannot be entirely attributed to loss of integrin binding^19^.

To determine if P66 is capable of binding to the CD47 receptor SIRPα, we developed an *in vitro* binding assay. Recombinant His-tagged P66 bound to immobilized SIRPα-Fc and CV1-G4 but not isotype control (Figure 1D and Extended Data Figure 1).

Utilizing a *p66*-deficient *B. burgdorferi* strain of B31-A3 (*Δp66*) we determined that P66 is required for CV1-G4 surface binding (Figure 2A). We next sought to determine residues on P66 critical for SIRPα interaction. We have previously demonstrated that two aspartic acid residues, D184 and D186, on a predicted extracellular loop of P66 (181-187) are required for integrin binding^19^. *B. burgdorferi* expressing the mutant D184A and D186A, *p66*^*D184A,D186A*^, or loss of the loop, *p66*^*Δ181-187*^, demonstrated loss of CV1-G4 binding (Figure 2A). Consistent to previous structure predictions, these sites map to an unstructured extracellular loop on a structure of P66 generated by Alphafold2 (Figure 2B and Extended Data 2A). We postulate this region is also required for SIRPα binding. Importantly, while these residues are critical for binding integrins and P66, loss of this loop or mutation of the two aspartic acid residues does not affect P66 cell surface localization^19^.

**Figure 2.**
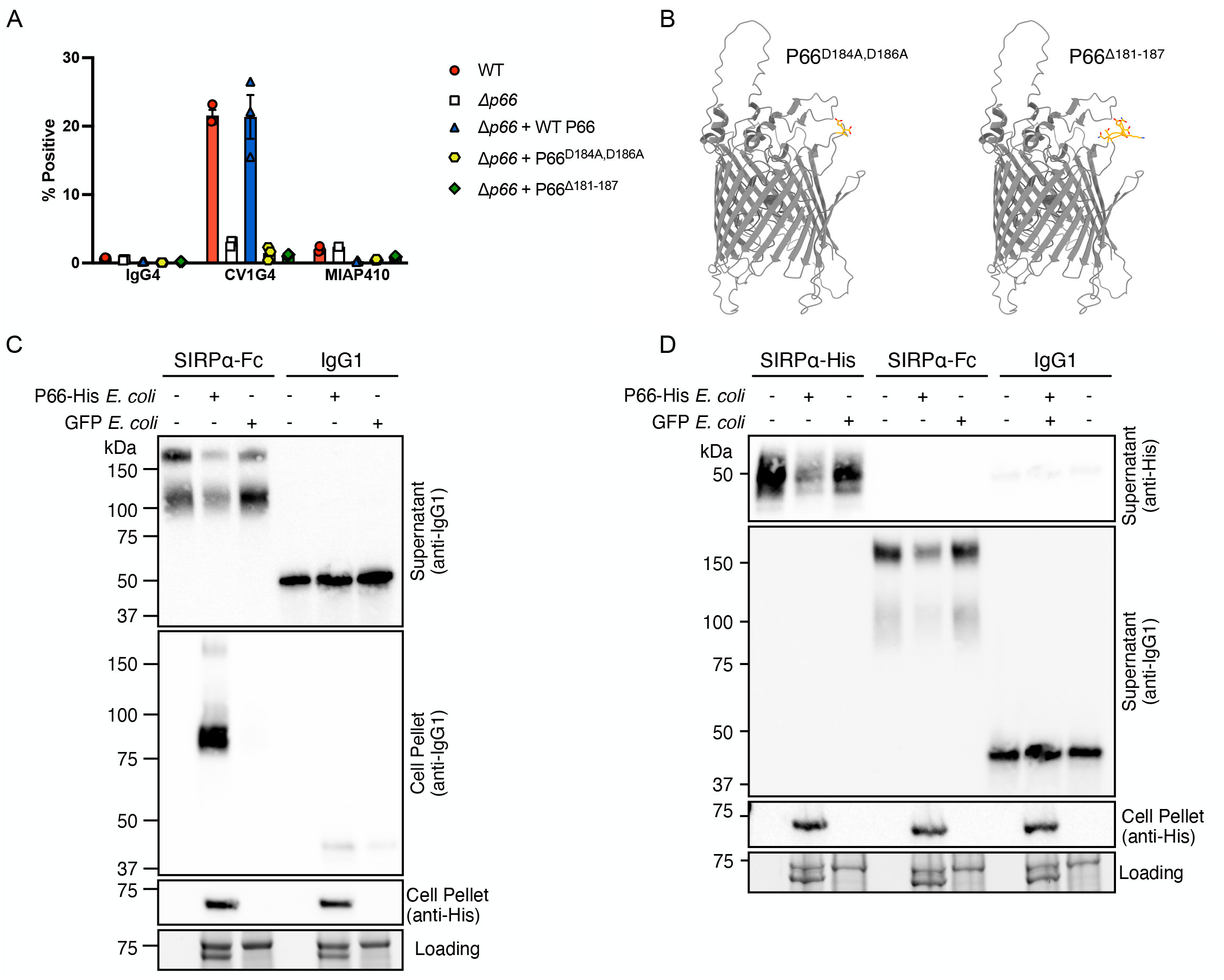
P66 is necessary and sufficient for binding to a CD47 affinity reagent and SIRPα. A. Wild-type (WT), *p66* deficient (Δ*p66*) *B. burgdorferi*, Δ*p66 B. burgdorferi* reconstituted with WT *p66*, Δ*p66 B. burgdorferi* reconstituted with the *p66*^*D184A D186A*^ mutant, or Δ*p66 B. burgdorferi* reconstituted with the *p66*^Δ*181-187*^ loop deletion mutant were stained with CV1-G4, IgG4 or MIAP410 and analyzed by flow cytometry. B. Predicted structure of P66 by Alphafold2 with mutated residues or loop deletions highlighted in orange. C. P66-His- or GFP-expressing *E. coli* were incubated with soluble SIRPa-Fc or IgG1 isotype control, prior to Western blot analysis. D. Soluble SIRPa-Fc, SIRPa-His or IgG1 isotype control was incubated with *E. coli* expressing P66-His or GFP prior to Western blot analysis.

We next asked whether expression of P66 is sufficient to bind SIRPα to bacteria. To this end we engineered *E. coli* to produce P66 on their cell surface and confirmed surface localization by detergent partitioning (Extended Data Figure 3). By pelleting assay^20^, we observed P66-dependent binding of SIRPα to the surface of *E. coli* and depletion of soluble SIRPα in the supernatant (Figure 2C and Extended Data Figure 4). To further validate that this depletion was dependent on the interaction between P66 and SIRPα, we subjected P66-expressing or GFP-expressing *E. coli* to incubation with recombinant purified SIRPα-Fc, the isotype control IgG4, or a his-tagged SIRPα which revealed P66-dependent depletion for both SIRPα reagents (Figure 2D and Extended Data Figure 5). Taken together these studies demonstrate that P66 is necessary and sufficient to bind to the human “don’t eat me” receptor SIRPα.

To investigate the role of P66 in mediating phagocytic clearance of *B. burgdorferi*, we performed phagocytosis assays using primary human monocyte-derived macrophages from 4 healthy donors challenged with WT or *Δp66 B. burgdorferi* strain B31-A3. WT and *Δp66 B. burgdorferi* were stained with a pH sensitive rhodamine dye to enable fluorescence imaging of *B. burgdorferi* localized to the acidic phagolysosome over the course of co-culture (158 h). Average counts of pHrodo-positive phagocytic events over time across all human macrophage donors co-incubated with *Δp66 B. burgdorferi* were higher compared to that of the WT *B. burgdorferi*, calculated through image segmentation on the fluorescence images (Figure 3A). Within the first 10 h of coincubation, *Δp66 B. burgdorferi* were phagocytosed at greater rates than WT (Figure 3B and 3C). Although there is variation between individual human macrophage donors in their peak event time and total number of events when challenged with *Δp66* or WT *B. burgdorferi* (Figure 3D), the ratio of pHrodo positive phagocytic events for *Δp66* compared to WT *B. burgdorferi* was similar across all donors (Figure 3E). Representative images and time lapse videos taken during the time of peak phagocytic events for WT and *Δp66 B. burgdorferi* co-culture and visually demonstrate the role of P66 in macrophage mediated phagocytosis (Extended Data Figures 6-8).

**Figure 3.**
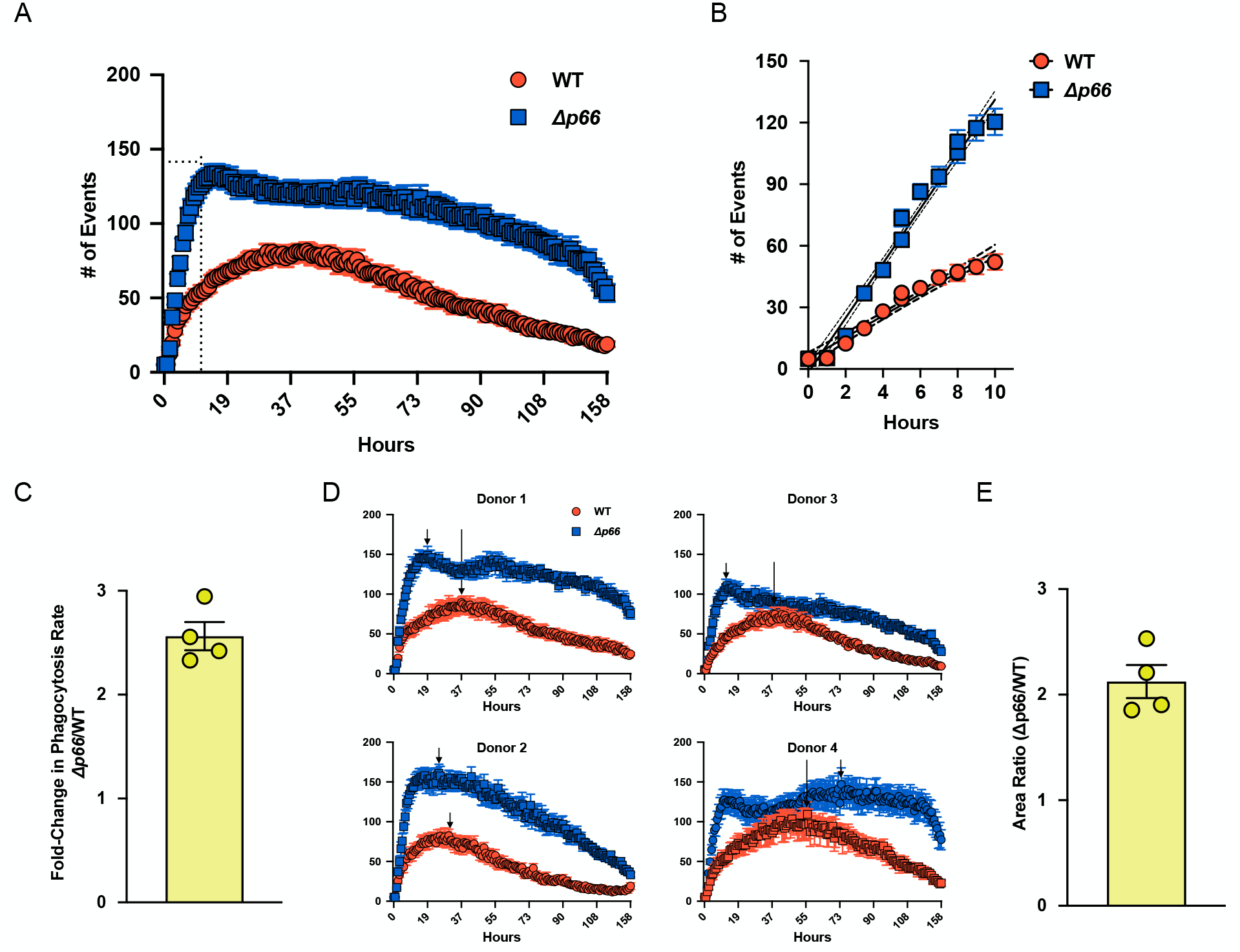
*Δp66 B. burgdorferi* are more-readily phagocytosed by human macrophages compared to WT as measured by pHrodo-positive phagocytic events. A. Phagocytosis assays using human monocyte derived macrophages averaged across 4 donors reveal that *Δp66* bacteria are more-readily taken up by macrophages throughout the course of the experiment. 2 technical replicates per donor with 3 fields of view per technical replicate, allowing for 6 fields of view measured per donor. B. *Δp66 B. burgdorferi* have a greater rate of increase in phagocytic events during the first 10 hours. 95% confidence intervals for the event rate are shown as dashed lines. C. Fold-change in rates of phagocytic events between *Δp66* and WT bacteria were calculated for each donor over the first 10 hours. D. Individual macrophage phagocytosis data from 4 human donors, 2 technical replicates per donor with 3 fields of view per technical replicate for a total of 6 measurements per donor. Arrows denote maximum for each donor with each strain. E. Fold-increase in total number of phagocytic events with *Δp66 B. burgdorferi* compared to WT across co-incubation time. Error bars calculated as SEM.

*B. burgdorferi* strain B31-A3 deficient in the gene encoding P66 are readily cleared from sites of infection on a time scale consistent with that of an innate immune response^18,19^. *In vivo* studies have demonstrated that CV1-G4 treatment promotes macrophage clearance of cancer cells in patient derived xenografts of bladder cancer and lymphoma in immunodeficient mice^14^. To further evaluate the therapeutic potential of P66 blockade in altering the course of infection, we infected mice with a derivative of *B. burgdorferi* strain ML23 expressing luciferase (ML23-pBBE22luc)^21,22^.

Importantly, this strain provides a readout of infection burden over the entire animal, allowing for the quantification of live bacterial load as well as dissemination. Mice were injected with 100,000 luciferase-expressing *B. burgdorferi* pre-treated with CV1-G4 or IgG4 isotype control. Live bacterial load and dissemination was determined by live animal imaging following injection with D-luciferin at Day 0, Day 7, and Day 14 (Figure 4A). Within the first day of infection, a significant reduction in radiance from bacteria pre-treated with CV1-G4 over isotype control was already observed. This trend continued at Day 7, suggesting that blockade of P66 can slow the kinetics of infection and impact the health of the bacteria. Notably, CV1-G4 is not toxic to *B. burgdorferi* in culture (data not shown). We wondered if infection with a larger bolus of bacteria could also be impacted with blockade of P66. Mice were injected with 1,000,000 luciferase-expressing *B. burgdorferi* pre-treated with CV1-G4 or IgG4 isotype control (Figure 4B). The average radiance was determined by live animal imaging at Day 0 and Day 9 following injection with D-luciferin, revealing a marked reduction in live bacteria between Day 0 and Day 9 with CV1-G4 pre-treatment.

**Figure 4.**
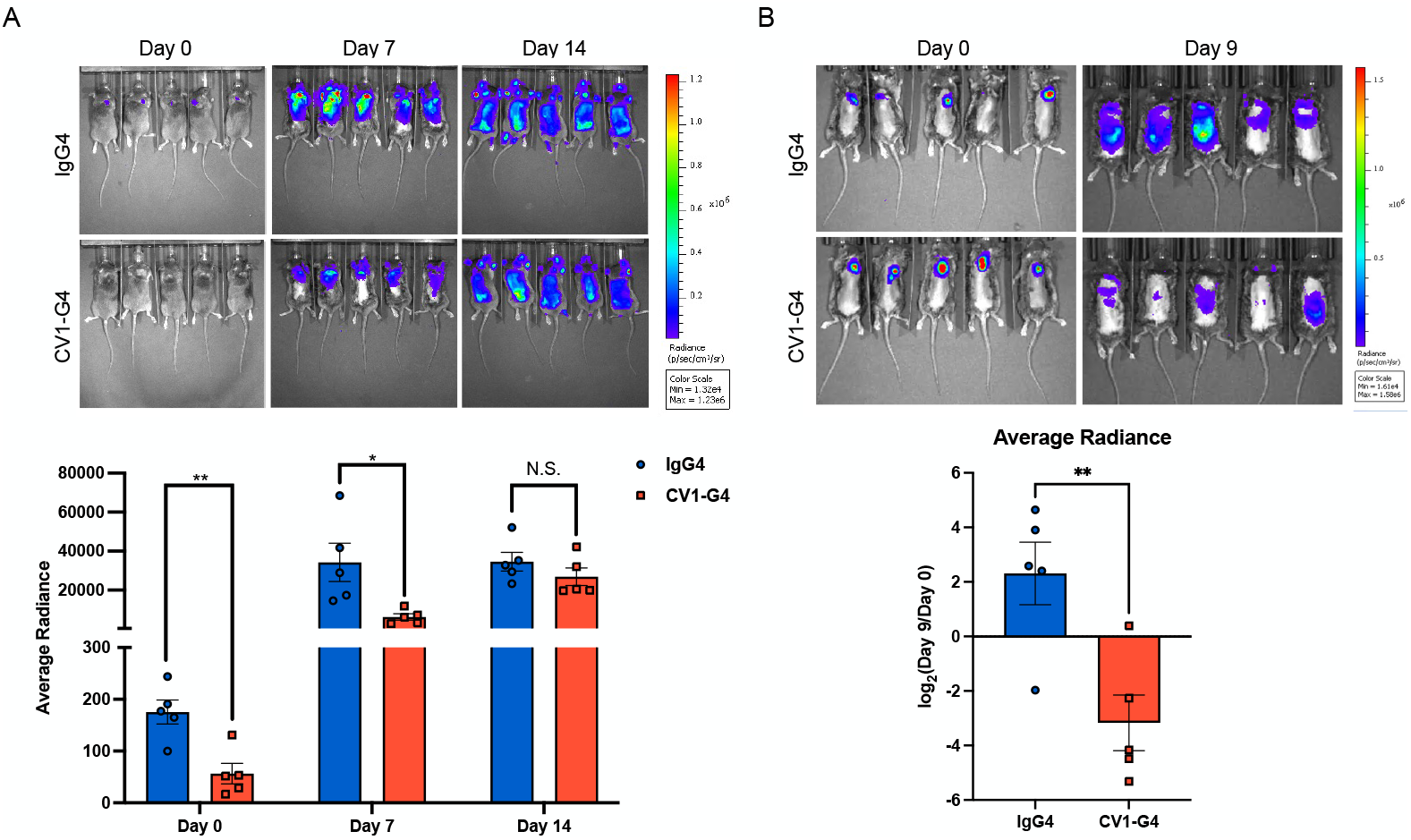
P66 is a therapeutic target in Lyme Disease. A. C3H Female mice were infected with 10^5^ luciferase-expressing *B. burgdorferi*, strain ML23-pBBE22luc, that were pretreated with CV1-G4 or isotype control (6 μg). Mice were imaged at Day 0, Day 7, and Day 14 following injection with D-luciferin for *in vivo* imaging. Error bars calculated as SEM. ** p-value = 0.00478; N = 5; * p-value = 0.04541; N = 5; N.S. = not significant. Unpaired Welch’s t-test. B. C3H Female mice were infected with 10^6^ luciferase-expressing *B. burgdorferi*, strain ML23-pBBE22luc, that were pretreated with CV1-G4 or isotype control (6 μg). Mice were imaged at Day 0 and Day 9 following injection with D-luciferin for *in vivo* imaging. Error bars calculated as SEM. Ratio of average radiance for Day 9/Day 0 was calculated for each mouse. ** p-value = 0.0075; N = 5; Unpaired Welch’s t-test.

To the best of our knowledge, these findings are the first validation of a mimic of a mammalian “don’t eat me” signal ligand encoded by pathogenic bacteria. We have demonstrated that *B. burgdorferi* express a CD47-like protein on their surface that promotes evasion of macrophage-mediated phagocytosis. We identified this protein as P66, a protein that has previously been implicated in persistent infection^15^. P66 is necessary and sufficient for SIRPα binding. Loss of *p66* increases *B. burgdorferi* clearance by human macrophages. Blockade of the P66-SIRPα interaction *in vivo* promotes bacterial clearance. These data suggest that P66 promotes *B. burgdorferi* evasion of innate immune cell-mediated clearance by phagocytes, functioning as a true “don’t eat me” signal ligand via SIRPα. Thus, we report P66 as a therapeutic target in the treatment of Lyme Disease.

Protein-wide, conformational similarity between P66 and CD47 is unlikely (Extended Data Figure 2). CD47 harbors an N-terminal Ig domain, extensive glycosylation sites, and 5 transmembrane domain sequences. None of these features are identified on P66; in fact, P66 is thought to form a beta-barrel and function as a porin^23,24^. However, both associate with integrins^11,16,17^. Alignment between *B. burgdorferi* P66 and human CD47 reveals no regions of significant sequence similarity. Given that humans are an accidental host for *B. burgdorferi* and the aforementioned evidence, we postulate that CD47 mimicry by *B. burgdorferi* is an instance of convergent evolution at the SIRPα binding site, rather than co-evolution. Future studies include solving the structure of P66 bound to SIRPα. P66 is encoded across the *Borrelia* genus, and therefore P66-like molecules may serve as a surface “don’t eat me” signal ligand for other *Borrelia* species. The broader applicability of this finding to other bacterial pathogens is of great interest to our laboratories and is currently being investigated.

CD47 is upregulated in cancer cells^19^, pathogen-infected host cells, and pathogenic cells in fibrosis^20,21^ and atherosclerosis^22^. Our finding of a role for P66 as a CD47 mimic in evasion of phagocyte-mediated immune clearance of *B. burgdorferi* parallels cancer’s use of the CD47-SIRPα checkpoint axis. While the field of immuno-oncology has investigated the therapeutic potential of targeting macrophages through immune checkpoint blockade, this blockade may also assist in the treatment of pathogenic infections that are notoriously challenging to clear, such as Lyme Disease. Currently, we rely exclusively on antibiotics for resolving bacterial infections, but these studies suggest that immunomodulatory targets may prove therapeutically synergistic. Doxycycline is a bacteriostatic antibiotic that halts the replication of bacteria. We postulate that if bacteria can evade phagocytic clearance, a course of bacteriostatic antibiotics may not always be sufficient to clear persistent bacterial infection^25^. Therefore, blockade of the P66-SIRPα checkpoint pathway may have potential as a supportive immunotherapy in the treatment of Lyme Disease.

## METHODS

### *B. burgdorferi* culture

In a sterile biosafety cabinet, stock vials of *B. burgdorferi* (1x10^7^ *B. burgdorferi*/tube), stored at -80ºC, were thawed into BSK-H complete medium with 6% rabbit serum (50 mL, Sigma-Aldrich) or BSK-II complete medium with 6% rabbit serum (50 mL, generously made by the Chou lab or made in-house following protocol from Jenny Hyde’s Lab). Strain *B. burgdorferi* B31 A3 GFP was generously provided by George Chaconas. Strains B31 A3 *p66* wildtype (WT) and B31 A3 *p66* KO C3-14 (*Δp66*) were generated as described previously and plasmids profiled prior to use in experimnts^18,19^. ML23-pBBE22luc was generously provided by Jenny Hyde^21,22^. For the *Δp66 B. burgdorferi* cultures, we selected for the mutation in kanamycin (200 μg/mL, Sigma-Aldrich). Each culture was cultured in sealed tubes at 37ºC unless otherwise stated.

### CV1-G4 and MIAP410 *B. burgdorferi* staining

*B. burgdorferi* cultures (GFP, wildtype or *Δp66*) (20 μL/well) were plated in a 96 well v-bottom plate (Corning). After the plate was centrifuged for 10 minutes at 1500 x g at 4ºC and the supernatant was aspirated, the *B. burgdorferi* were resuspended in FACS buffer (30 μL/well) (200 μL; 2% fetal bovine serum in PBS supplemented with EDTA (1 mM, Thermo Fisher Scientific)) containing CV1-G4 or MIAP410 (10 μg/mL). Samples were incubated on ice (30 min) and protected from light. Upon incubation completion, the plate was washed twice with PBS (150 μL). For each wash, bacteria were pelleted prior to aspiration (1500 x g, 10 min, 4ºC). After the second wash, the *B. burgdorferi* were resuspended PBS (30 μL) containing Alexa Fluor 647 anti-human IgG (1:200, Jackson ImmunoResearch) or Alexa Fluor 647 anti-mouse IgG (1:200, Jackson ImmunoResearch). Samples were incubated on ice (30 min) and protected from light. Samples were again washed with PBS (2x) as described above. *B. burgdorferi* were resuspended in 4% paraformaldehyde (100 μL, EMS) and incubated while protected from light (10 min, 25 ºC). Samples were again washed with PBS (2x) as described above prior to resuspension in FACS buffer. Samples were protected from light until analysis on a BD Fortessa.

### Fluorescence-Activated Cell Sorting (FACS)

The concentration, viability of cells and percentage of GFP expression was calculated with Fluorescence-Activated Cell Sorting (FACS). At Stanford University School of Medicine FACS Core, FACS was conducted on a BD LSRFortessa with BD FACS Diva software. For FACS analysis of *B. burgdorferi* at Stanford, LSRFortessa cytometer threshold levels were modified to parameter SSC 400 and voltages were set to FSC 300 and SSC 230, collected in log mode. At Massachusetts Institute of Technology Koch Institute Flow Cytometry Core, FACS was conducted on either a BD Symphony A3 HTS I or a BD LSRFortessa HTS II. For FACS analysis of *B. burgdorferi* at MIT, Symphony cytometer threshold levels were modified to parameter SSC 400 and voltages were set to FSC 250 and SSC 240, collected in log mode. For FACS analysis of *B. burgdorferi* at MIT, LSRFortessa cytometer threshold levels were modified to SSC 800 and voltages were set to FSC 300 and SSC 240. For all FACS assays, *B. burgdorferi* were counted and assessed for integrity but not assessed for motility. For all IncuCyte assays, *B. burgdorferi* were counted and assessed for integrity via FACS and assessed for motility via IncuCyte imaging.

### Enrichment of CV1-G4 candidate binding proteins

GFP-expressing *B. burgdorferi* (strain B31-A3, gifted by George Chaconas) was grown according to culture procedures and tested for CV1-G4 staining by FACS as described above. To generate cultures that stain negative for CV1-G4, bacteria were grown at 25ºC. Bacteria (150 mL) was then pelleted by centrifugation (500 x g, 10 min) and growth media aspirated away. Pellet was resuspended in ice-cold PBS and re-pelleted (500 x g, 10 min). This process was repeated an additional 2 times to ensure no residual growth serum from the media remains. The pellet was then resuspended in 1 mL of PBS, transferred to a 1.5 mL tube, and re-pelleted a final time (500 x g, 10 min). Non-denaturing lysis buffer (150 μL, 1% Triton X-100, 50 mM Tris-HCl, pH 8.5, 150 mM NaCl) containing protease and phosphatase inhibitors (EDTA-free Protease Inhibitors and Phos-STOP, Roche) was added and the pellet resuspended by pipetting and brief bath sonication during which time lysate should solubilize. Lysate was incubated on ice (15 min) and protein concentration determined by BCA Assay (Pierce ThermoFisher Scientific). Protein concentration was normalized to 1 μg/μL using additional ice-cold lysis buffer for a total of 300 μg.

CV1-G4 (1.5 μL, 1:200 dilution, 5 μg/μL stock) or an isotype control (IgG4, 7.5 μL, 1:40 dilution, 1 μg/μL) was added to the lysate from *B. burgdorferi* known to express a CV1-G4 binding protein or known to not express as previously confirmed by FACS. The lysate and reagents were allowed to incubate on a rotator (1.5 h, 4 °C). Prior to incubation completion Pierce Protein A/G Agarose Beads (ThermoFisher Scientific, 30 μL per 1 mg protein) were washed with ice-cold PBS (1 mL) and pelleted by centrifugation (1600 x g, 2 min, 4 °C). Washed beads were then resuspended in 5x volume of ice-cold lysis buffer and transferred to the antibody/reagent-incubated cell lysate samples. Samples were then incubated to enrich for antibody/reagent bound proteins of interest (1.5 h, 4 °C). Upon incubation completion, samples were moved to ice and pelleted by centrifugation (1600 x g, 2 min, 4 °C). The supernatant was aspirated away and beads washed with ice-cold lysis buffer (1 mL) followed by pelleting (1600 x g, 2 min, 4 °C). This was repeated 2x for a total of 3 washes. After final wash and supernatant removal, 1x Loading Buffer (40 μL, 3:1 Lysis Buffer:4x Laemmli Loading Buffer (Bio-Rad)) was added to the sample and the sample boiled (10 min, 95 °C) to elute bound samples.

Sample was loaded onto an AnyKD Mini-PROTEAN gel (20 μL, Bio-Rad) and separated by SDS-PAGE. Gel was stained with Colloidal Blue Staining Kit according to manufacturer’s protocol (ThermoFisher Scientific). Following detstaining with water, gel bands of interest were excised and subjected to preparation for mass spectrometry analysis.

### Preparation of gel bands for mass spectrometry analysis

Gel bands were diced into approximately 1 mm cubes and transferred to a 1.5 mL tube. Add 50 mM 1:1 50 mM NH_4_HCO_3_:Acetonitrile (200 μL) and shake (250 rpm, 30 min). Using a gel-loading tip (VWR) buffer was removed, which was blue as colloidal blue was removed. This was repeated 1x, after which time gel bits were clear. Gel bits were dehydrated by addition of acetonitrile (100 μL) and became opaque white. Acetonitrile was then removed and the gel bits were allowed to air dry in a fume hood (5 min).

Gel bits were rehydrated with reducing buffer (30 μL, 10 mM dithiothreitol in 50 mM NH_4_HCO_3_) and incubated with shaking (30 min, 45 °C). Excess buffer was removed with a gel-loading tip and replaced with alkylation buffer (30 μL, 55 mM iodoacetamide in 50 mM NH_4_HCO_3_). Sample was allowed to incubate (45 min, 25 °C) prior to removal of excess buffer. Gel pieces were then incubated with 50 mM NH_4_HCO_3_ (200 μL, 10 min with shaking) prior to dehydration with acetonitrile (200 μL). Acetonitrile was then removed, and opaque gel bits dried in a fume hood (10 min). Trypsin was reconstituted in 50 mM acetic acid in water at a concentration of 1 μg/μL (Trypsin Gold, Promega). Reconstituted trypsin was added to 50 mM NH_4_HCO_3_ in a 1:50 ratio (v/v). The resulting buffer was added to dehydrated gel bits (15 μL) and the gel bits sat at room temperature for 10 min before determined to be rehydrated by visual inspection. Upon rehydration, additional 50 mM NH_4_HCO_3_ was added as needed to ensure gel slices were completely submerged in buffer. Samples were allowed to incubate with shaking (16 h, 37 °C).

Upon incubation completion, samples were acidified with 1% TFA (5 μL), vortexed and allowed to rest (10 min). Gel bits were pelleted by centrifugation and supernatant collected using a gel-loading tip. Extraction buffer was added to each sample (50 μL, 60% acetonitrile, 1% TFA) and allowed to sonicate in a bath sonicator (10 min). Upon sonication completion, gel bits were pelleted, and supernatant was collected with a gel loading tip. This extraction process was repeated 1x. Collected sample was concentrated to dryness by SpeedVac. Samples were resuspended in 2% Acetonitrile, 0.1% Formic Acid prior to mass spectrometry analysis.

### Mass spectrometry data acquisition and search

In a typical mass spectrometry experiment, an Orbitrap Q Exactive HF-X mass spectrometer (Thermo Scientific, San Jose, CA) coupled to an Acquity M-Class UPLC system (Waters Corporation, Milford, MA) was used for separation and analysis. Reverse-phase separations were performed on a ∼25 cm fused silica chromatography column that was pulled-and-packed in-house. This fused silica column had an O.D. of 360 microns and an I.D. of 100 microns, and was packed under higher than operating pressure (∼12000 psi) with a C18 reprosil Pur 1.8 micron stationary phase (Dr. Maisch). The mobile phases were 0.2% aqueous formic acid and 0.2% formic acid in acetonitrile for phase A and B, respectively. Reconstituted peptides were directly injected onto the chromatography column, with a gradient of 2-45% mobile phase B, followed by a high-B wash over a total 80 minutes. The mass spectrometer was operated in a data-dependent mode using Higher Energy Collisional Dissociation (HCD) fragmentation for MS/MS spectral generation detected in the Orbitrap.

Mass spectra were analyzed using Byonic v 1.4.0 (Protein Metrics, Cupertino, CA) against the *B. burgdorferi* strain B31 database on Uniprot, including containing common contaminants and PTMs, *e*.*g*. methionine oxidation. Precursor and HCD fragment mass tolerances were set to 12 ppm. Data were validated using the standard reverse-decoy technique at a 1% false discovery rate as described previously^26^.

### Recombinant P66 Binding Assay

Recombinant his-tagged P66 was purchased from ProspecBio in a solution of 20mM HEPES. The protein was diluted to a final concentration of 2.5ng/μL in 200μL (500ng per sample) of Buffer A (20mM HEPES, 7.5 in MilliQ water). CV1-G4, isotype control, or SIRPα-Fc (5ng/μL final concentration, 1ug) was added to the diluted P66 protein and rotated at 4°C for 1.5 hrs. Protein G magnetic affinity beads were prepared with 3x200μL washes of buffer A. 2 μL of beads were prepared for each sample. Washed beads were suspended in 10x volume of buffer A (20μL) to ease transfer to samples. Beads were transferred to samples containing the mixture of P66 with CV1-G4, isotype control, or SIRPα-Fc at 4 °C for 1.5 hrs. Samples were rinsed 3x with buffer A before resuspension in 10μL 2x SDS Loading buffer with 350mM BME and 5μL buffer A. Samples were boiled (5 min., 95°C). The entire sample was loaded onto a TGX Stain Free Precast Gel (Biorad) in 1x Tris/Gly buffer and run at 200V for approximately 45 mins. The gel was then imaged for loading using the stain-free method on a ChemiDoc XRS+ (Biorad). Gel was incubated overnight at 4°C in transfer buffer (50mM Tris, 40mM Glycine, 0.1% SDS, 20% MeOH).

Following incubation protein was transferred from gel onto PVDF membrane. Blot was blocked with 5% milk in 1xTBST (1 h, 25°C). Blot was then incubated with 1:1000 dilution anti-his conjugated to HRP (Thermo Scientific) in 5% milk in 1xTBST (1 h, 25°C). Blot was then washed 3x with 1xTBST, (5 min, 25°C). Following washes, blot was incubated in Radiance Plus ECL (Azure Biosystems) for approximately 5 minutes. Blot was visualized using the chemiluminescent setting on a ChemiDoc XRS+ (Biorad).

### Bacterial strains and growth conditions

*E. coli* were grown at 30 °C in Luria-Bertani (LB) broth (VWR).

### Cloning and expression of GFP and recombinant P66

Our starting P66 construct was gifted from Melisha Kenedy, who cloned P66 from *B. burgdorferi* B31 genomic DNA into pET23a, which contains a C-terminal 6x-His tag, as described^27^. Using NEBuilder HiFi DNA assembly (NEB), the *E. coli* signal peptide for OmpT (ATGCGGGCGAAACTTCTGGGAATAGTCCTGACAACCCCTATTGCGATCAGCTCTTTTGCT) was inserted into Kenedy’s P66 plasmid. Codon optimization of the OmpT-P66 sequence for expression in *E. coli*, and secondary expression in human, was performed by GenScript. The GFP plasmid (pZE27GFP) was a gift from James Collins (Addgene plasmid 75452). Both the P66 and GFP constructs were transformed into the *E. coli* strain BL21(DE3)pLysS (Promega).

For expression of P66, 25 mL of LB broth was inoculated with 500 μL of an overnight culture. Cultures were grown at 30 °C to an optical density at 600 nm (OD_600_) of 0.2, then P66 expression was induced with 1 mM isopropyl-β-D-thiogalactopyranoside (IPTG) and grown at 18 °C overnight to a final OD_600_ of 0.9-1.2. For expression of GFP, 25 mL of LB broth was inoculated with 250 μL of an overnight culture and cultures were grown at 30 °C to an OD_600_ of 0.9-1.2.

### Expression and purification of SIRPα with a C-terminal 6x-His tag (SIRPα-His)

25 mL of Expi293F cells were transfected following the manufacturer’s recommended protocol. The transfected plasmid contained SIRPA (NM_001040022), a.a. 1-349 followed by a C-terminal 6x-His tag in a pcDNA3.1 backbone. 18 hours after transfection, enhancers were added and the flask was moved to 30 °C. After 6 days at 30 °C, the flask was harvested and the media was filtered with a 0.2 μm filter. Pierce High-Capacity Ni-IMAC resin was equilibrated with equilibration buffer (50 mM NaH_2_PO_3_, 300 mM NaCl, 10 mM imidazole, pH 8) before incubation with the filtered media for 1 hour at 4 °C in batch. Flow through was collected using a gravity column and the resin was washed 3 times with equilibration buffer + 20 mM imidazole. The desired protein was eluted in 0.5 mL fractions with elution buffer (50 mM NaH_2_PO_3_, 300 mM NaCl, 500 mM imidazole, pH 8). Fractions containing the desired protein were combined and dialyzed overnight in 50 mM NaH_2_PO_4_, 300 mM NaCl, pH 8. Dialysis was repeated in fresh buffer. Protein concentration was measured by BCA assay.

### Triton X-114 phase partitioning

To study the amphiphilic properties of P66, detergent phase partitioning was performed as described previously^28^. Aqueous- and detergent-enriched proteins were analyzed by SDS-PAGE and immunoblotting. For SDS-PAGE analysis, protein pellets were resuspended in 50 μL of PBS (phosphate-buffered saline, pH 7.4) then mixed with 4x Laemmli sample buffer (Biorad), boiled for 5 min at 95°C, and loaded onto Criterion TGX stain-free protein gels (Biorad). After electrophoresis, stain-free gels were imaged and then transferred onto polyvinylidene fluoride (PVDF) membranes for immunoblot analysis. Blots were blocked for 30 min to 1 h in 5% milk in TBST (1X Tris-buffered saline with 0.1% Tween 20) and then incubated with an anti-6X-His tag primary antibody (1:1,000 dilution, Invitrogen) (overnight, 4°C). After 3x washes with TBST (5 min, 25°C), blots were incubated with anti-mouse IgG, HRP-linked secondary antibody (1:1,000 dilution, Cell Signaling Technology) (1 h, 25°C). The blot was washed 3x with TBST again, incubated in Radiance ECL HRP substrate (Azure Biosystems) for 2 min and visualized using the chemiluminescent setting on a ChemiDoc XRS+ (Biorad).

### Bacterial pelleting assay

To analyze sufficiency of P66 for binding SIRPα, a bacterial pelleting assay adapted from Martin et al^20^, was performed. 10^9^ *E. coli* expressing P66, or GFP as a negative control, were suspended in 200 μL of a 125 nM solution of recombinant human SIRPα-Fc protein (R&D Systems), the cognate isotype control recombinant human IgG1 protein (Sino Biological), or SIRPα-His. Samples were then incubated at 30 °C for 30 min while shaking (220 rpm). Cultures were centrifuged (500 rcf, 5 min) and the supernatant was transferred. Cell pellets were washed twice with 450 μL of PBS. Supernatant and cell pellet samples were analyzed with SDS-PAGE and immunoblotting. For SDS-PAGE analysis, supernatant samples were mixed with 4x Laemmli sample buffer. *E. coli* cell pellets were snap frozen in liquid nitrogen, then resuspended in 75 μL of PBS and 25 μL of 4x Laemmli sample buffer and lysed in an ultrasonic bath for 1 min. All samples were loaded onto Criterion TGX stain-free protein gels (Biorad) for gel electrophoresis. The stain-free gel was imaged and transferred onto a PVDF membrane. The blot was blocked for 30 min to 1 h in 5% milk in TBST. For immunoblotting, an anti-human IgG, Fcγ-specific, HRP-linked antibody (1:5,000, Cell Signaling Technology) was used for detection of SIRPα-Fc and IgG1. P66 and SIRPα-His were detected using an anti-6x-His tag, HRP-linked antibody (1:500 dilution, Invitrogen). Blots were incubated with antibodies overnight (4°C), then 3x washes with TBST (5 min, 25°C) and incubated in Radiance ECL HRP substrate (Azure Biosystems) for 2 min before visualization using the chemiluminescent setting on the ChemiDoc XRS+ (Biorad).

### Primary human donor-derived macrophage generation and stimulation

Leukocyte reduction system chambers from anonymous healthy donors were obtained from the Stanford Blood Center. Peripheral monocytes were purified through successive density gradients using Ficoll (Sigma-Aldrich) and Percoll (GE Healthcare). Monocytes were then differentiated into macrophages by 7–9 days of culture in IMDM with glutamax base + 10% AB human serum (Life Technologies).

### Time-lapse live-cell-microscopy-based phagocytosis assay

Human serum derived macrophages were lifted using TrypLE (5 mL, Thermo Fisher Scientific). Once lifted, the macrophages in TrypLE were diluted with R10 without phenol red (5 mL) (10% Fetal Bovine Serum in RPMI without phenol red (Thermo Fisher Scientific)). The macrophages were counted using Trypan Blue (Life Technologies) and centrifuged (300 x g, 5 min, 4ºC). After aspirating the supernatant, the macrophages were resuspended in R10 without phenol red (1.5x10^5^ cells/mL). The macrophage suspension was plated in each well (100 μL, 150,000 macrophages/well) of a 96-well Imagelock plate (Essen). The plate was incubated to ensure macrophage adherence (37ºC, 30 min). *Δp66* and WT *B. burgdorferi* B31-A3 cultures were centrifuged (1500 x g, 10 min, 4ºC) and resuspended into pHrodo (1:20,000, Essen) in PBS (10 mL). After incubation (1 hour, 37ºC), fetal bovine serum (1 mL) was added to each *B. burgdorferi* suspension. The *B. burgdorferi* were pelleted and resuspended in R10 without phenol red (3x106 *B. burgdorferi*/mL). After the macrophages adhered to the Imagelock plate, the *B. burgdorferi* solution (50 μL, 1,500,000 *B. burgdorferi* per well) was plated into each well at an MOI of 10.R10 without phenol red (100 μL) was added to wells without macrophages and R10 without phenol red (50 μL) was added to wells without *B. burgdorferi*. Phagocytosis assay plates were placed in an IncuCyte (Essen) within an incubator (37 °C) and imaged at 45-60-min intervals for 6.57 days. The first image time point (reported as t = 0) was acquired within 30 min of co-culture. Images were acquired using a 20x objective at 800-ms exposures per field. *B. burgdorferi* were assessed for motility via IncuCyte imaging. Phagocytosis events were calculated as the number of pHrodo-red+ events per fields of view. Threshold values to determine pHrodo-red+ events were set such that macrophages not incubated with pHrodo labeled *B. Burgdorferi* had no pHrodo-red+ events.

### Murine *B. burgdorferi-*luciferase infection bioluminescent quantification

ML23-pBBE22luc *B. burgdorgeri* strains (kindly gifted by Dr. Jenny A. Hyde) were cultured for 4 days at 37ºC in BSK-II media and complete with 6% rabbit serum (Millipore Sigma) at pH of 7.6. ML23-pBBE22luc cultures had the addition of a selection antibiotic, kanamycin at 300 μg/mL (Sigma-Aldrich). Bacterial concentration was determined by flow cytometry (Becton Dickinson LSR-II). 1,000,000 spirochetes per condition were co-incubated with IgG4 (Biolegend) or CV1-G4 (6 μg) for 1 hour at 37ºC. The culture was then resuspended in 500 μL of 0.2% uninfected B6 murine serum (diluted in PBS). Female C3H/HeJ mice (n=5 per condition) aged 10 weeks were provided by Jackson Laboratories (Bar Harbor, ME). Mice were anaesthetized by isoflurane inhalation and injected intradermally with 100,000 *B. burgdorferi* in 50 μL. To confirm infection, an injection of 100 μL sterile filtered D-luciferin (GoldBio) reconstituted in sterile PBS at 277 mg/kg was done intraperitoneally. The hair on the backs of the mice was removed using an electric shaver, then the addition of Nair for 30 seconds to 1 minute. Gauze wipes were used to clean the backs in the following order: soaked in 70% ethanol, soaked distilled water, and a dry gauze. After 15 minutes, mice were arranged in the In Vivo Imaging System (Perkin Elmer) then imaged for an exposure time of 1 minute. This in vivo imaging using D-luciferin was later repeated at each of the indicated intervals (days 0, 7 or 9, 14).

### Analysis of In vivo Imaging System (IVIS)

All downstream IVIS analyses were performed blinded in Living Image (Perkin Elmer). The average radiance values (photons/sec/cm^2^/sr) were achieved as the sum of radiance from each pixel within the rectangular gating of individual mice, extending from the top of the nose cone to the tail tip. All images were normalized across all cages and time points with a set radiance range from 1.55e4 -1.48e6 (p/sec/cm2/sr) for visualization only.

### Protein Alignment

Protein alignments were performed through Uniprot (www.uniprot.org) using the Clustal Omega Program^23,24^. The following proteins were used for analysis: CD47_HUMAN (Q08722), and H7C7N8(P66)_BORBU (H7C7N8).

### Statistical Analysis

All statistical methods and number of replicates are reported in figure legends. Error bars were calculated as standard error to the mean (SEM) in GraphPad Prism.

## Acknowledgements

We thank members of the Weissman, Zaro and Robinson laboratories and the Stanford Stem Cell Institute for advice and discussions; T. Naik and L. Quinn for technical and logistical support. We thank N. Ramadoss, B. Hayes, Y-Y. Yiu, R. Brewer, O. Colace, J. Ross, A. Ring, J.-P. Volkmer, and R. Salomon for contributions of lab reagents and enabling work. We thank Jenny Hyde for the gift of the ML23-pBBE22luc strain of *B. burgdorferi* and George Chaconas for the B31-A3 GFP strain of *B. burgdorferi*. The research reported in this publication was supported by the Emily and Malcolm Fairbairn Donor-Advised Fund (to M.C.T., I.L.W., and B.W.Z.); The Bay Area Lyme Foundation (M.C.T., I.L.W., and B.W.Z.); the Robert J. Kleberg, Jr., and Helen C. Kleberg Foundation (to I.L.W.); the Virginia D. K. Ludwig Fund for Cancer Research (to I.L.W.); Arnold and Mabel Beckman Foundation (Beckman Young Investigator to B.W.Z.); the Program in Breakthrough Biomedical Research, partially funded by the Sandler Foundation (to B.W.Z.); Life Sciences Research Foundation fellowship from the SVCF-Wave Fund (to A.L.S.); Stanford Immunology training grant 5T32AI007290 (to M.C.T.); the NIH NRSA 1 F32 AI124558-01 award (to M.C.T.); The Lyme Disease Association (Grant Award #: A140241, to E.E.B.), the Deutsche Forschungsgemeinschaft (DFG, German Research Foundation) under Germany’s Excellence Strategy – EXC 2124–390838134 (to E.E.B.).This work was supported by the Vincent Coates Foundation Mass Spectrometry Laboratory, Stanford University Mass Spectrometry (RRID:SCR_017801) utilizing the Thermo QE-HFX mass spectrometer (RRID:SCR_018703). This work was supported in part by NIH P30 CA124435 utilizing the Stanford Cancer Institute Proteomics/Mass Spectrometry Shared Resource. The funders had no role in study design, data collection and analysis, decision to publish, or preparation of the manuscript.

## Author Information

### Contributions

M.C.T. conceived and designed experiments, performed experiments, analyzed data, edited the manuscript, and supervised the research. P.S.H., H.A.O., Q.F., and R.F.V. performed experiments, analyzed data, and wrote the manuscript. B.L. performed experiments. S.L.C. performed experiments and edited the manuscript. M.S. and S.G. performed experiments. A.L.S. provided reagents and edited the manuscript. L.B.T.D. performed experiments and edited the manuscript. V.R.P., H.H.P., and J.R. provided reagents. E.E.B. provided expertise and edited the manuscript. S.C. provided reagents and edited the manuscript. J.C. provided the *p66* mutant and WT parental *B. burgdorferi* strains, provided advice, and edited the manuscript. W.H.R. provided expertise and edited the manuscript. I.L.W. conceived and designed experiments, supervised the research and edited the manuscript. B.W.Z. conceived and designed experiments, performed mass spectrometry experiments, analyzed data, supervised the research, and wrote the manuscript.

### Ethics declarations

M.C.T., W.H.R., B.W.Z. and I.L.W. are co-inventors on a patent application (63/107,295) related to this work. I.L.W. is an inventor on U.S. patent 2019/0092873 A1 CD47, Targeted Therapies for the Treatment of Infectious Disease. I.L.W. was cofounder, director, and stockholder in FortySeven Inc., a public company that was involved in CD47-based immunotherapy of cancer during this study but was acquired by Gilead. At the time of this submission, I.L.W. has no formal relationship with Gilead. S.C. is currently a board member for Holoclara and co-founder and CEO of Arcadia Science. Her contribution to this work was completed before these affiliations. Animal studies were performed at the Department of Comparative (DCM) at the Massachusetts Institute of Technology (MIT) (Cambridge, MA). All procedures and care guidelines were approved by the MIT Committee on Animal Care (CAC) (Protocol#1221-087-24).

---

**Extended Data Figure 1.**
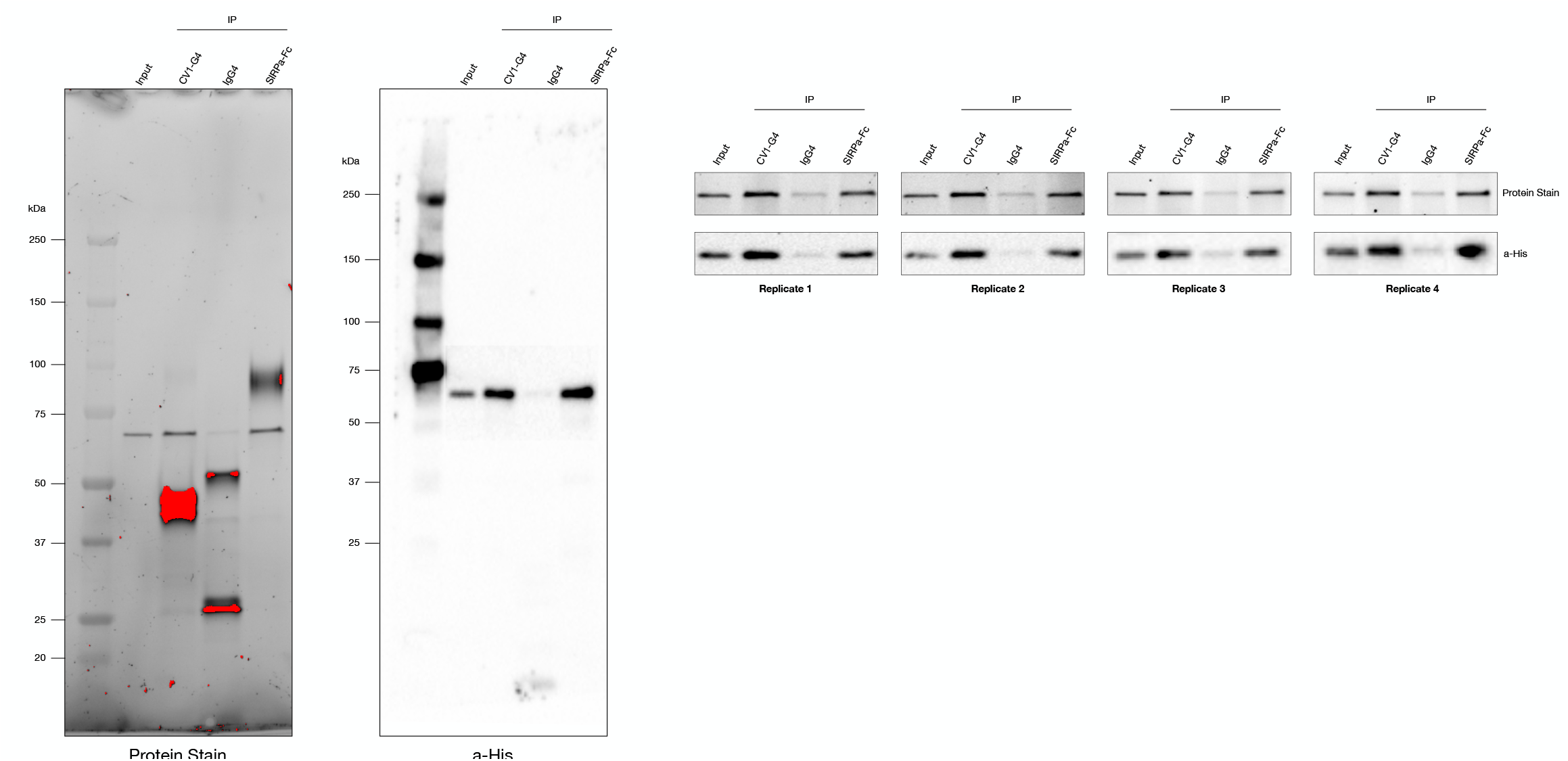
Recombinant P66 is enriched by CV1-G4 or SIRPα but not isotype control as determined by *in vitro* binding assay. Representative full gel and Western blotting images of all 4 replicates of binding assay quantified in Figure 1D.

**Extended Data Figure 2.**
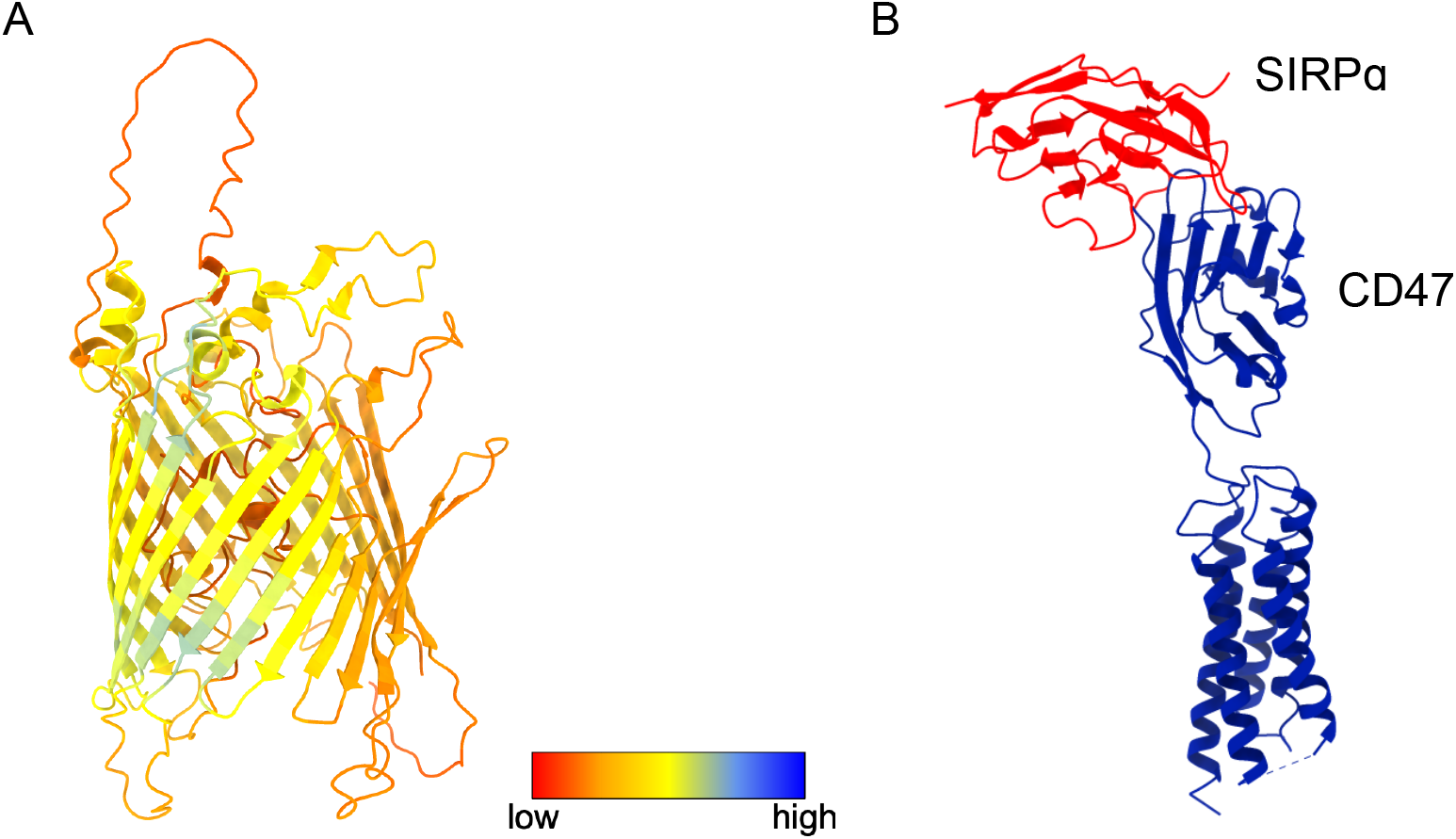
P66 and CD47 are structurally distinct proteins. A. Structure of P66 as predicted by Alphafold2. B. The human CD47-SIRPα binding interface. The crystal structure of the IgSF domain of CD47 bound to the N-terminal IgV domain SIRPα (PDB: 2JJT) was aligned with the cryo-EM structure of full-length human CD47 (PDB: 7MYZ).

**Extended Data Figure 3.**
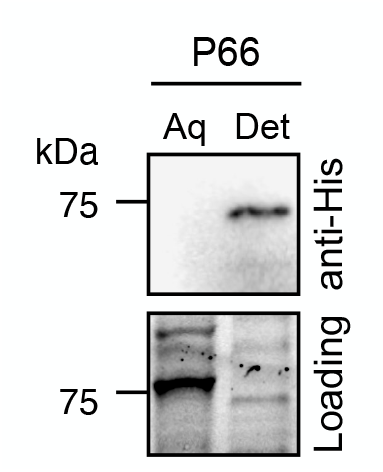
Detergent partitioning of P66-expressing *E. coli* reveals P66 is selectively detected in the membrane (detergent) fraction by anti-His Western blot.

**Extended Data Figure 4.**
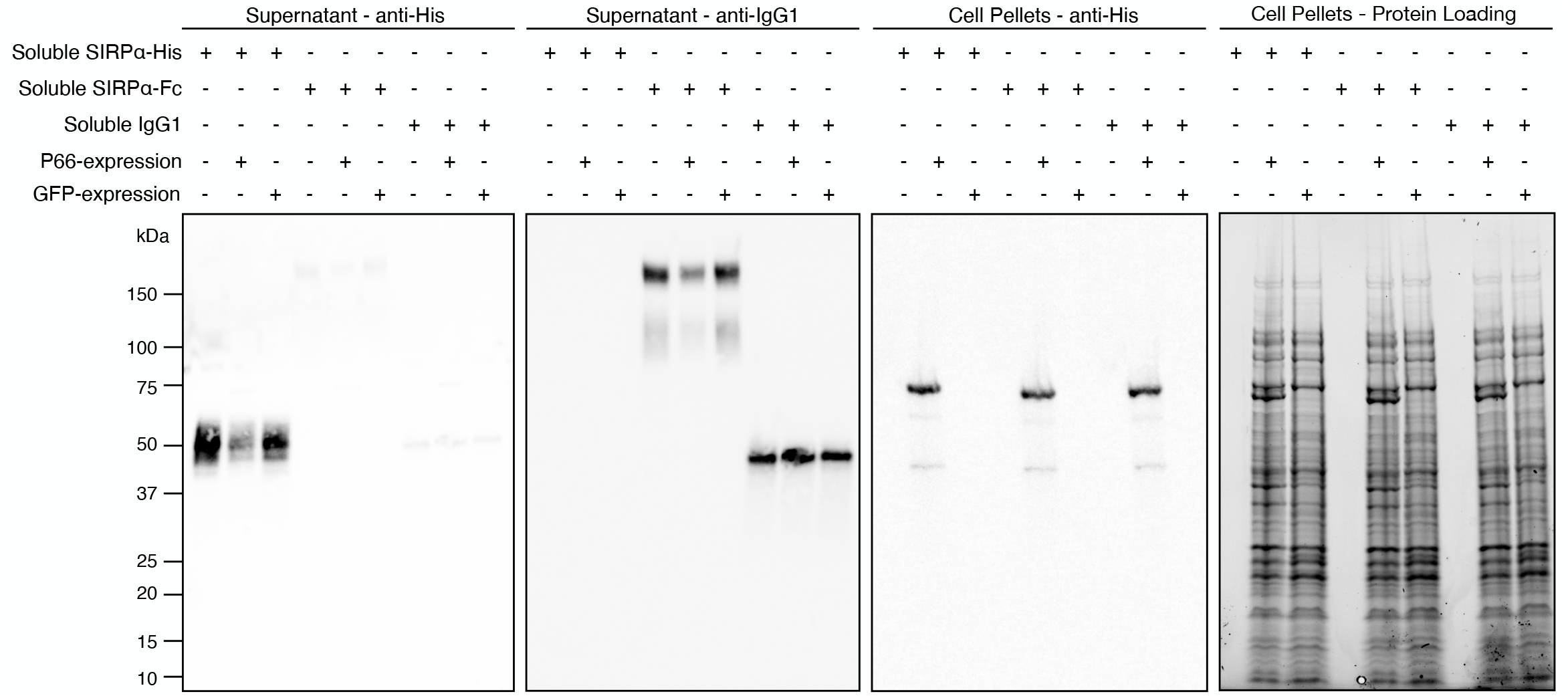
Full Western blot and gels from pelleting assay described in Figure 1C. P66- or GFP-expressing *E. coli* were incubated with soluble SIRPa-Fc or isotype control, prior to Western blot analysis.

**Extended Data Figure 5.**
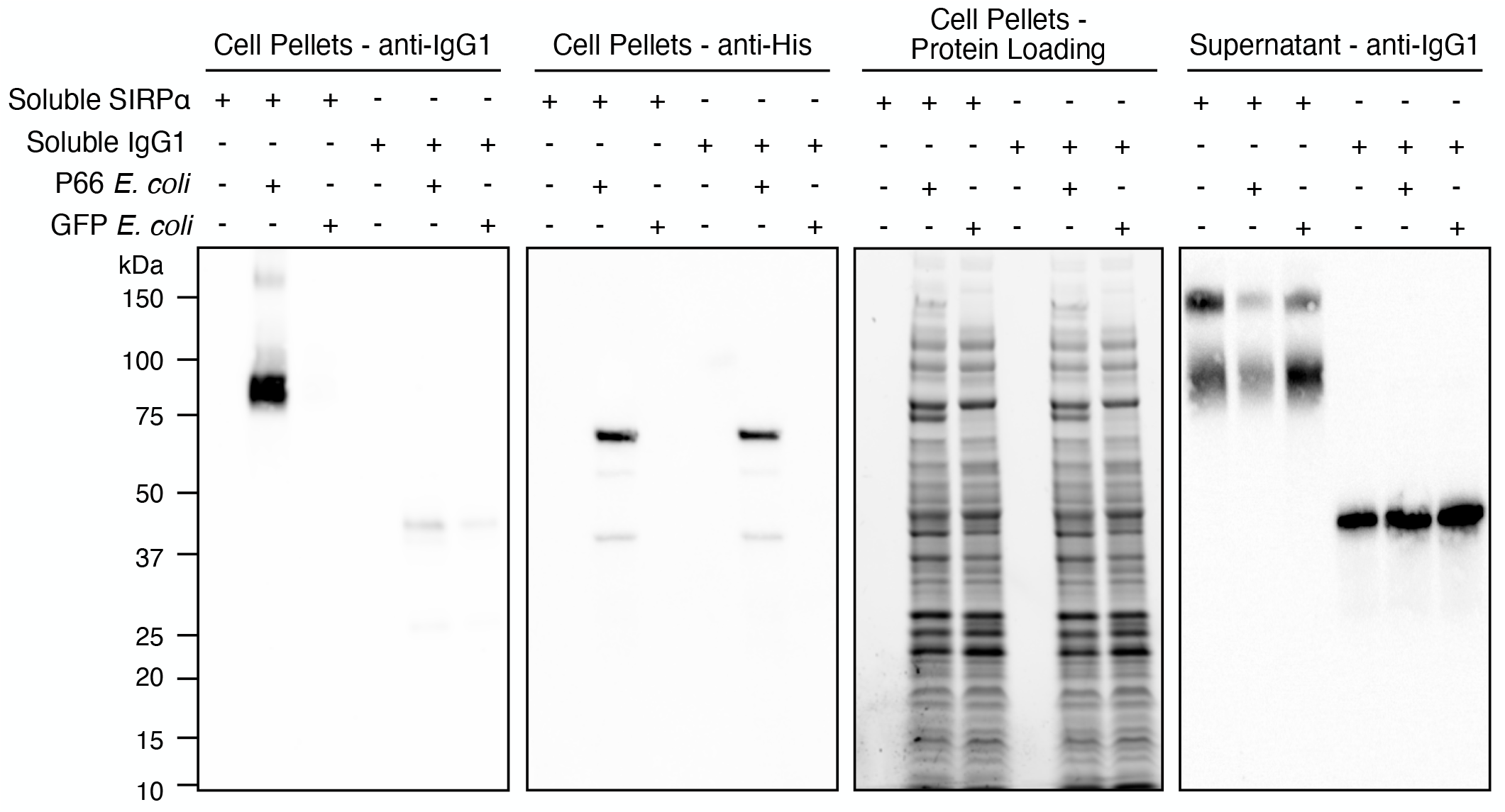
Full Western blot and gels from cell pelleting assays described in Figure 1D. Soluble SIRPa-Fc, SIRPa-His or isotype control was incubated with *E. coli* expressing P66 or GFP prior to Western blot analysis.

**Extended Data Figure 6.**
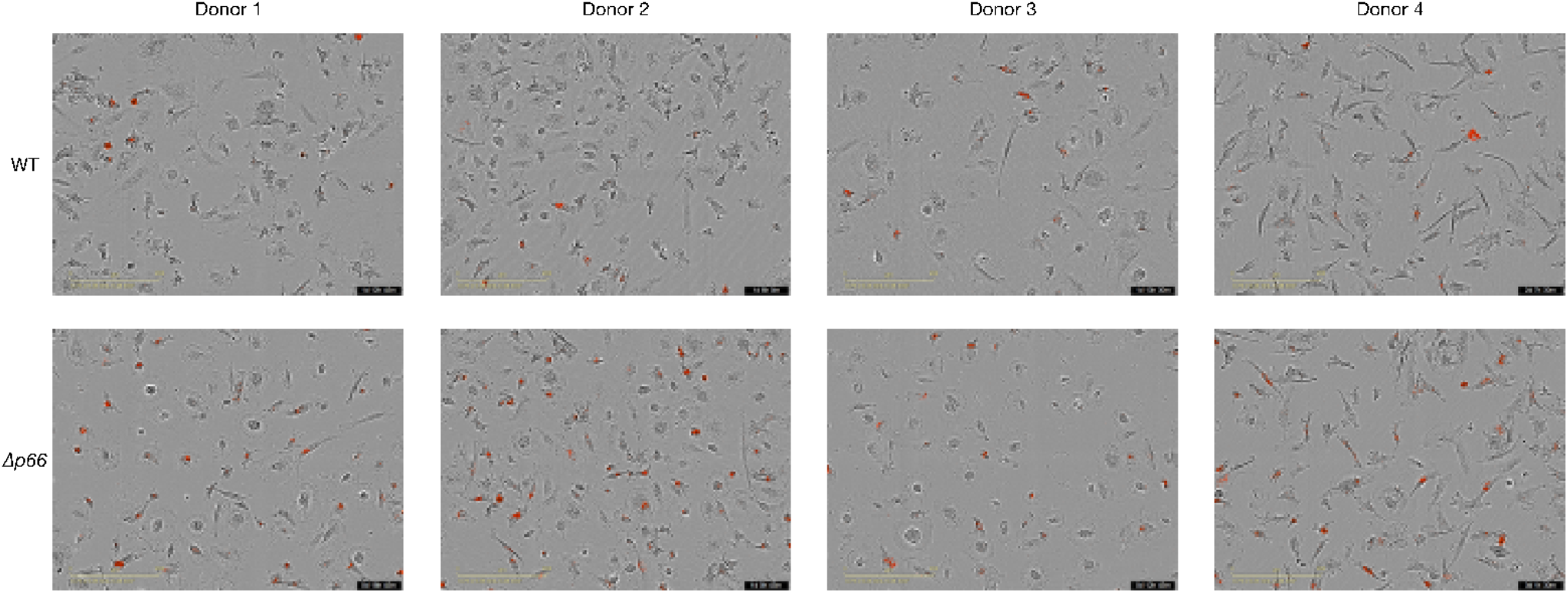
Peak phagocytosis images with macrophages from 4 donors (denoted by arrows in Figure 3D). Red events indicate lysosomal localization of wild-type (WT) or *p66* knockout (*Δp66*) *B. burgdorferi*.

<u>WT phagocytosis video</u>

**Extended Data Figure 7.** Time-lapse videos of phagocytosis of wile-type B31-A3 *B. burgdorferi* (WT) by macrophages. Red events indicate lysosomal localization of *B. burgdorferi*. Hosted on Google Drive.

*<u>p66</u>*<u>knockout phagocytosis video</u>

**Extended Data Figure 8.** Time-lapse videos of phagocytosis of *p66* knockout B31-A3 *B. burgdorferi* by macrophages. Red events indicate lysosomal localization of *B. burgdorferi*. Hosted on Google Drive.

